# How to make “number lines” stable in the mind’s eye

**DOI:** 10.1101/714659

**Authors:** Mario Pinto, Michele Pellegrino, Fabio Marson, Stefano Lasaponara, Vincenzo Cestari, Fabrizio Doricchi

## Abstract

Humans are prone to mentally organise the ascending series of integers according to reading habits, so that in western cultures small numbers are positioned to the left of larger ones on a mental number line (MNL, Dehaene et al., 1993). Despite 140 years since seminal observations made by Sir Francis Galton in human observers that experienced vivid MNLs (Galton, 1880a,b), the neural and functional mechanisms that give rise to MNLs remain elusive. Here, in a series of three experiments we show that spatially organised MNLs are not “all or none” phenomena and that MNLs get progressively less noisy and more stable in the mind’s eye, the more an observer fully combines left/right spatial codes with small/large magnitude codes to operate on numbers. These results show that conceptualising the ascending series of integers in spatial terms depends on contextual clues rather than being inherent to the semantic representation of numbers.

## Introduction

Getting acquainted with reading and writing in one’s own culture powerfully shapes the ability of conceptualising numbers in spatial terms. Several studies have suggested that the mental representation of the ascending series of integers runs from left-to-right in left-to-right reading cultures and vice versa in right-to-left reading ones (Dehaene, Bossini, & Giraux, 1993; Shaki, Fischer, & Petrusic, 2009). This way of representing number magnitudes, the so called “Mental Number Line” (MNL), is especially marked when reading habits are the same for words and numbers, like in western cultures or in Palestinians, while it can be less pronounced in cultures, like in Israeli Hebrew speakers, where reading directions for words and numbers run opposite (Shaki, Fischer, & Petrusic, 2009). MNLs also do not strictly depend on visual experience, as they occur both in sighted participants and in blind people who have acquired reading direction through the manual modality (for review see Bottini et al., 2015). One crucial issue in the study of the functional origins of MNLs is establishing how deep is the link that reading habits create between space and numbers. Do, as an example, the learned associations between “left” and “small numbers” and between “right” and “large numbers” go so deep that the concept “left” will inherently activate the concept “small number” and vice versa? Is the generation and use of spatially oriented MNLs rooted in these types of co-activations?

## Method

To give an answer to these questions, we have tested 84 right-handed adult healthy participants that were assigned randomly to three different groups of 28 participants each (statistical power > .90 with α set to .05; see SOM). Each group performed a different version of a task (Experiments 1-3) that by recording the timing of simple unimanual responses to intermixed numerical and spatially-informative visual targets, provides a measure of the mental association between numbers and space (Nosek and Banaji, 2001; Fischer and Shaki, 2017). In different trials of this task, an Arabic digit ranging from “1” to “9” or an arrow that points to the left or to the right, acts as target in the centre of the visual field (see S1 in SOM). The presence of an active MNL that runs from left to right, is revealed by faster responses when the direction of task-relevant arrow-targets is congruent with the position that task-relevant number-targets would occupy on the MNL, e.g. “push when an arrow points left and when a number is small “, rather than when it is incongruent, e.g. “push when an arrow points right and when a number is small” (Nosek and Banaji, 2001; Fischer and Shaki, 2017). Here we tested the significance and stability of this congruency effect in task conditions that, over three different experiments, promoted progressively the full conceptual integration of small/large codes of numerical-magnitude with left/right spatial codes of arrow direction.

## Results

In Experiment 1, participants were asked to respond to targets using rules that required the discrimination of the left/right direction of arrows without requiring the discrimination of the small/large numerical magnitude of digits or, vice versa, the discrimination of the small/large numerical magnitude of digits without requiring the discrimination of the left/right direction of arrows. Therefore, in different blocks of trials participants had to respond (Figure 1A): a) only to left pointing arrows and to all digits independently of their magnitude; b) only to right pointing arrows and to all digits; c) only to digits lower than 5 and to all arrows independently of their left/right direction; d) only to digits lower than 5 and to all arrows. In this experiment responding only to left pointing arrows did not speed up the detection of digits lower than 5, which would have been expected if a shared conceptual representation of “left” and “small” was active, and responding only to right pointing arrows did not speed up the detection of digits higher than 5 [Space-to-Number influence: t (27) = 1.83, p = .10, d = .37; average = 4.26, SD = 11.38; Exp. 1, Figure 1B; split-half reliability: r_1,2_ = -.24, r_tt_ = .39, p = .25, see bottom panel Figure 1B]. Using the same logic, responding only to digits lower than 5 did not speed up the detection of arrows that pointed to the left, i.e. toward the position that these digits should occupy on the MNL, and responding only to digits higher than 5 did not speed up the detection of arrows pointing to the right [Number-to-Space influence: t (27) = .26, p = .79, d = .05; average = .96, SD = 17.82 Exp.1, Figure 1C; split-half reliability: r_1,2_ = -.36, r_tt_ = .53, p = .10, see bottom panel Figure 1C]. Taken together, these results show that MNLs are not evoked when left/right spatial concepts are contrasted without contrasting also small/large magnitude codes and vice-versa.

**Figure 1.**
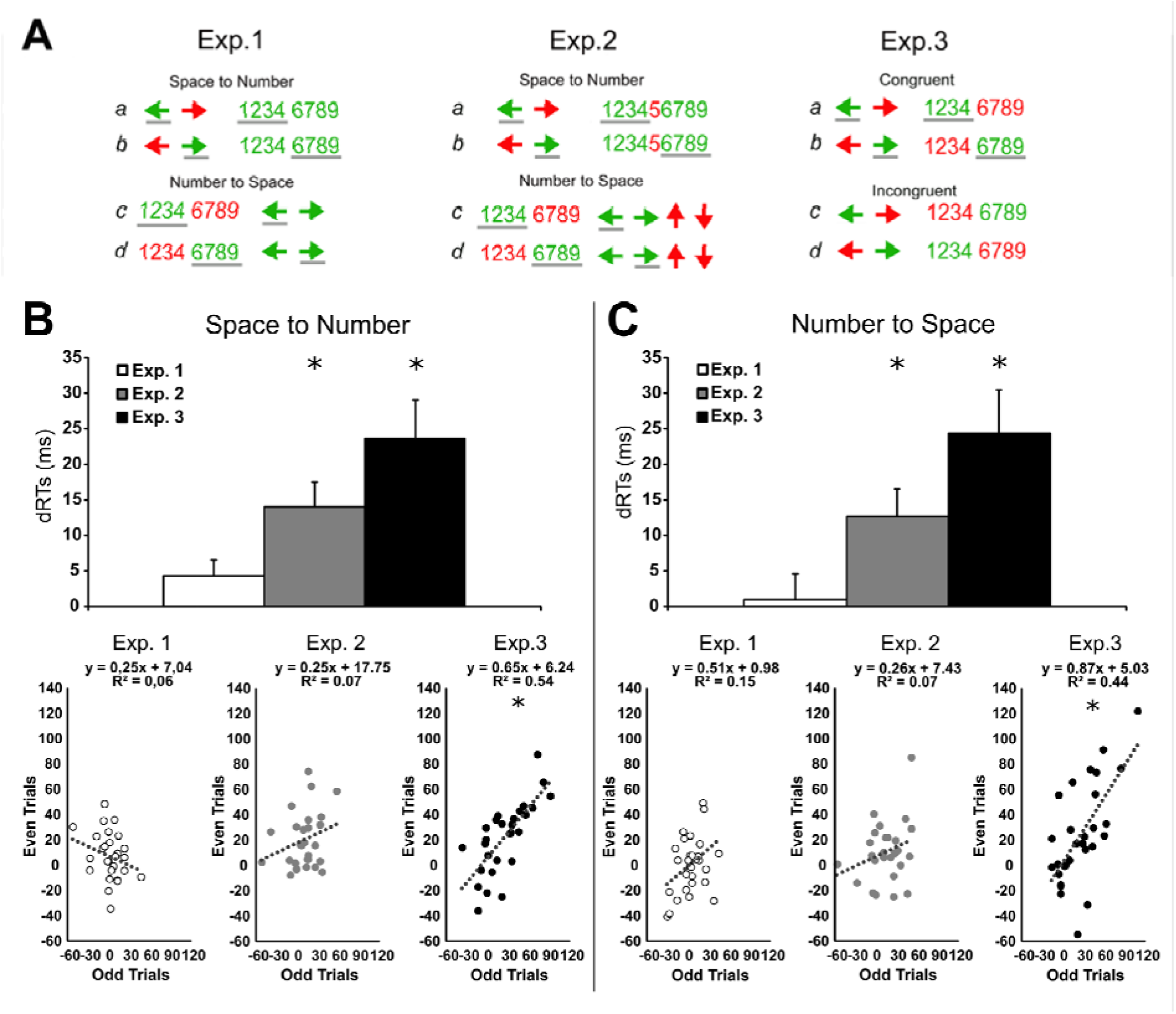
**A)** Go targets (in green) and NoGo targets (in red) in the different experimental conditions of Experiment 1,2 and 3. Participants had to perform unimanual speeded key presses only to Go targets while refraining from responding to NoGo targets. Grey lines highlight Go spatial and numerical targets for which faster detection should be expected if a conceptual association between spatial codes, i.e. Left/Right, and number magnitude codes, i.e. Small/Large, is active during task performance. ***B) Top panel: Space-to-Number influence*** = Reaction Time advantage (dRTs) in the detection of numerical targets when participants are also engaged in the detection of directionally congruent arrows (Left Arrow/Small Magnitude digits, Right Arrow/Large Magnitude digits) with respect to the detection of directionally incongruent arrows (Left Arrow/Large Magnitude digits, Right Arrow/ Small Magnitude digits); ***Bottom panel:*** scatterplots representing the stability of the Space-to-Number influence in Experiment 1,2 and 3 (Split-half reliability); ***C) Top panel: Number-to-Space influence*** = Reaction Time advantage (dRTs) in the detection of left and right pointing Arrows when participants are also engaged in the detection of numerical targets that occupy directionally congruent position on the MNL (Small Magnitude digits/ Left Arrow, Large Magnitude digits/ Right Arrow) with respect to the detection of numerical targets in incongruent position on the MNL (Small Magnitude digits/Right Arrow, Large Magnitude digits/Left Arrow); ***Bottom panel:*** scatterplots representing the stability of Number-to-Space influence in Experiment 1,2 and 3 (Split-half reliability). Thin bars in histograms represent SE. Statistically significant effects are marked with an asterisk (see Main text and SOM for details).

In Experiment 2, we wished to explore to which degree each of the left/right and small/large conceptual dichotomies must be explicitly called into action to produce MNLs when both spatial and numerical-magnitude codes are relevant to the performance of the task. In Experiment 2 participants used both contrasting left/right spatial codes for the discrimination of arrow direction and contrasting small/large codes for the discrimination digit magnitude though, in different experimental conditions, the two codes included in one of these contrasts were explicitly activated by task instructions, while the two codes included in the remaining contrast were only implicitly activated through superordinate semantic knowledge: i.e. “*horizontal*” is superordinate to “left/right” and “*different from* 5” is superordinate to “smaller/larger than 5”. Therefore, in different blocks of trials participants had to respond (Figure 1A): a) only to left pointing arrows and to *digits different from 5;* b) only to right pointing arrows and to *digits different from 5*; c) only to numbers lower than 5 and to *horizontal arrows* (that over different trials were intermixed with vertical arrows that pointed up or down); d) only to numbers higher than 5 and to *horizontal arrows*. In this task, attending to arrows in a specific left or right direction speed-up the detection of directionally congruent magnitudes on the MNL, i.e. small and large respectively, though no small/large discrimination of magnitude was explicitly required by the task [t (27) = 4.30, p < .001, d = .81; average = 14.09, SD = 17.33; Exp.2 Figure 1B]. In a similar way, attending to a specific small or large digit magnitude speed-up the detection of directionally congruent left-pointing or right-pointing horizontal arrows respectively, though no discrimination of left/right arrow direction was explicitly required [t (27) = 3.47, p < .01, d = .66; average = 12.66, SD = 19.29; Exp.2 Figure 1C]. Nonetheless, reliability tests showed that these interactions between space and numbers were unstable and not consistent across different trials during task performance [Space-to-Number: r_1,2_ = .26, r_tt_ = .41, p = .18, bottom panel Figure 1B; Number-to-Space: r_1,2_ = .27, r_tt_ = .43, p = .16, bottom panel Figure 1C].

In Experiment 3, we finally tested whether significant and stable MNLs are generated when participants are asked to discriminate arrow-direction and number-magnitude basing on instructions that fully and explicitly combine left/right spatial with small/large magnitude codes. In four different experimental conditions participants had to respond (Figure 1A): a) only to arrows pointing left and to digits smaller than 5; b) only to arrows pointing right and to digits larger than 5; c) only to arrows pointing left and to digits larger than 5; d) only to arrows pointing right and to digits lower than 5. In this case, significant associations between space and numbers were found, as responding on the basis of instructions that explicitly promoted the association between conceptually congruent spatial and magnitude codes, i.e. “small” with “left” and “right” with “large”, speed-up the discrimination of both arrow direction and digit magnitude compared to instructions that were based on conceptually incongruent pairings, i.e. “left” with “large” and “right” with “small” [Space-to-Number: t (27) = 4.47, p < .001, d = .84; average = 23.71, SD = 28.05; Exp.3 Figure 1B; Number-to-Space: t (27) = 4.04, p < .001, d = .76; average = 24.96, SD = 32.67; Exp.3 Figure 1C]. Most important, these space-number associations were not only significant but also fully stable during task performance [Space-to-Number: r_1,2_ = .73, r_tt_ = .85, p <. 001, bottom panel Figure1B; Number-to-Space: r_1,2_ = .66, r_tt_ = .78, p < .001, bottom panel Figure 1C; see SOM]. Space-number congruency was significantly higher and more stable in Exp. 3 than in Exp. 1 and Exp. 2 (all p < .01; see SOM).

## Discussion

Although mathematical discoveries often defy and mismatch visuospatial expectations that are shaped by sensory experience, visual thinking has an indisputable and polymorphous role in the making, teaching and understanding of mathematics (Giaquinto, 2007). MNLs, in particular, provide a simple tool for grasping the meaning and representing integers, rational, real, complex numbers and the coordinate systems of planes and volumes. The results of our study open a window on the mechanisms that subtend the representational reliability and stability of space-number associations, because they show that the more left/right spatial codes and small/large magnitude codes are jointly and fully contrasted during the processing of numbers the more MNLs get noiseless and stable in the mind’s eye. This is particularly pointed out by the results of Exp.2, where the superordinate activation of spatial left/right or magnitude small/large code pairings was not sufficient to generate stable and reliable MNLs. Future studies should assess whether these findings also extend to the spatial conceptualization of other important cognitive dimensions as, for example, time (the “Mental Time Line”; Casasanto & Bottini, 2014; Bottini et al., 2015). The results of our study also suggest that the semantic representation of number magnitude has not an inherent spatial component (Aiello et al., 2012; Aiello et al., 2013; Harvey et al., 2013; Fattorini et al., 2015; Rotondaro et al., 2015; Pinto et al., 2018; Rosetti et al., 2011), as the isolated discrimination between “small” and “large” numerical magnitudes produced no activation of corresponding “left” and “right” spatial codes, respectively, and vice versa (Exp.1). Therefore, in line with recent high-resolution fMRI mapping studies that show no overlap between the topographical distributions in the cerebral cortex of neuronal populations that are sensitive to different positions in space and neuronal populations that are sensitive to different numerosities (Harvey et al., 2013), our data support a degree of independency between the brain representation of space and that of numbers. This functional independency might provide important developmental advantages as it would equally favour the acquisition of left-to-right or right-to-left organised MNLs depending on the cultural-context. In contrast, assuming a phylogenetically original and inherent left-to-right organization of the MNL (Rugani et al., 2015) would imply an onerous developmental re-organization of the space-number interaction in individuals that belong to right-to-left reading cultures who organize their MNL from right-to-left. Finally, the relative independency between the representation of numbers and that of space, fits well with the rich diversity of mathematical thinking, where both propositional-symbolic and visual-spatial reasoning have played relevant epistemic roles and have independently produced important advances (Giaquinto, 2007).

## Author contributions

F.D, M.P. and M.P developed the study concept. All authors contributed to the study design. M.P., M.P., F.M. and S.L. collected and analysed data. F.D., M.Pinto and V.C. drafted the manuscript. All co-authors provided critical revisions and rewrote the manuscript.

## Acknowledgements

We wish to thank Sarah Shomstein, Daniele Nico and Roberto Bottini for useful suggestions on a first draft of the manuscript. We thank Giada Baldo, Jessica Callà, Valentina Galiotta, Claudia Golser and Alessandra Martucci for help in data collection. This work was supported by grant PRIN 2017 (project code 2017XBJN4F) and grant Ricerche di Ateneo 2018 to F.D.

