## Supplemental File for "How to make “number lines” stable in the mind’s eye"

**1. General Method**
*1.1 Participants* Based on average effect sizes obtained by Shaki and Fischer (2018, Experiment 2), we determined the number of participants that would have been needed to obtain a power of .90 with alpha set to .05 (two-sided). A total sample size of 25 would be needed in each experiment (average Cohen's d = 0.68). We tested a total of 84 right-handed adult healthy participants that were assigned randomly to three different groups of 28 participants each

*1.2 Apparatus*
All experiments were run in a sound attenuated room with dim illumination. Stimuli were presented on a 15-inch-color VGA monitor. An IBM-compatible PC running MATLAB software controlled the presentation of stimuli and the recording of responses. Participants had their head positioned on a chin rest at a viewing distance of 57.7 cm from the screen. All participants had normal or corrected to normal vision and were naive to the experimental hypothesis of the experiment. Different samples of participants were included in the three experiments of the present study.

*1.3 Statistical analyses*
In each experiment, the significance of *Space-to-Number* (Congruent: Left Arrow /Small Number & Right Arrow /Large Number, Incongruent: Right Arrow /Small Number & Left Arrow /Large Number) and *Number-to-Space* (Congruent: Small Number/ Left Arrow & Large Number/ Right Arrow, Incongruent: Small Number/ Right Arrow & Large Number/ Left Arrow) congruency effects was evaluated by first calculating the individual RTs advantages produced in the Congruent with respect to the Incongruent Condition (dRTs = RTs in the Incongruent Condition *minus* RTs in the Congruent Condition), and then by entering these values in a series of t-test analyses.
 In a second step, the reliability of Space-to-Number and of Number-to-Space congruency effects observed in each experiment was assessed using the split-half method. We first calculated the RTs advantages produced in the Congruent with respect to the Incongruent Condition for the odd-numbered items and in the even-numbered items. Then, the corrected Spearman-Brown correlation between the RTs advantages in the odd-numbered and even-numbered halves of trials was used as index of reliability.
 To compare the size of Space-to-Number and Number-to-Space congruency effects among the three different experiments, we performed two series of null-hypothesis-significance-testing analyses (NHST). In both cases, we compared dRTs through one-way between experiments ANOVA.
 We also compared the reliability coefficients observed in the three different experiments. In a first step, for each experiments we re-calculated reliability coefficients by resampling data through the replacement of a different participant (n-1, bootstrap). Then we entered these coefficients in a one-way between experiments ANOVA.

Finally, we tested whether variability in reliability measures was comparable and homogeneous among the three different experiments. To this aim, for each experiment we first calculated individual ΔRTs of congruency effects (congruency effect in odd-numbered items *minus* congruency effect in even-numbered trials) and then entered these values in a one-way between experiments ANOVA. Go trials in which no response was provided (misses) and trails with RTs above 1000 ms or below 100 ms were not included in the analyses.

*1.4 Procedure*

Each trial started with the 500 ms presentation of a central fixation cross (1.5° × 1.5°). At the end of this delay an Arabic digit (1, 2, 3, 4, 6, 7, 8 or 9; size = 1.5° × 1°; font = Arial) or an arrow pointing to left or to the right (size = 1.5° × 0.8°), replaced the central fixation cross. Target stimuli remained available for response for 2000 ms. The inter-trial interval was 500 ms (see Supplementary Figure 1). The task was divided into 4 blocks of trials, each corresponding to a different experimental condition. All experiments (see detailed instructions in the main text) included four experimental conditions that were administered during a single experimental session. The order of experimental conditions was counterbalanced among participants. A short break was allowed between blocks. At the beginning of the experimental session, participants performed a short training session (16 trials).

*1.4.1 Experiment 1*
Each block consisted of 256 trials, 128 with numerical-targets and 128 with arrow-targets (64 trials per each of the two arrow-directions).

*1.4.2 Experiment 2*

The procedure of Experiment 2 was similar to Experiment 1, except that in this case also vertical arrows pointing up or down and the digit “5” were presented. Each block included 216 trials. In experimental conditions “a” and “b” 144 trials had numerical-targets (72 No-Go trials for digit “5”) and 72 arrow-targets (i.e. 36 trials for each arrow-direction). In experimental conditions “c” and “d” 72 trials had numerical-targets (note that in these conditions number “5” is not presented) and 144 arrow-targets (i.e. 72 trials for the Go horizontal arrow-direction and 72 trials for the No-Go vertical arrow-direction).

*1.4.3 Experiment 3*

Stimuli, procedure, number of blocks and order of experimental conditions, were the same of Experiment 1. Task instructions are described in the main text.

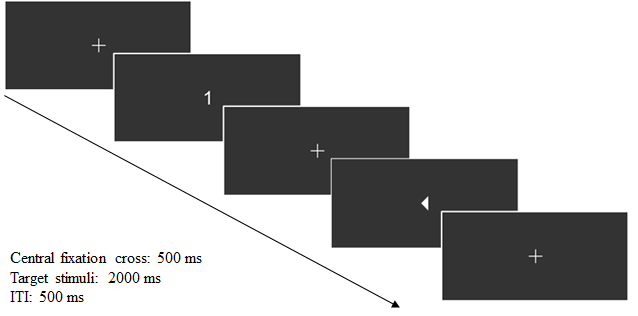

**Supplementary Figure 1**: *Examples of two consecutive trials, one with a numerical-target (1) and one with an arrow-target (arrow pointing to the left in this case).*

**1.5 Additional Results**
*1.5.1 Null-Hypothesis-Significance-Testing (NHST).*

To compare the Space-to-Number and Number-to-Space congruency effects that were observed in the three different experiments, we run two series of null-hypothesis-significance-testing analyses (NHST) using one-way between experiments ANOVAs.
As regards with Space-to-Number congruency, the ANOVA highlighted a significant main effect of Experiment [F (2, 81)

= 5.83, p

<

.01, η_p_^2^ = .13]. Post-hoc tests showed that in Exp.3 congruency (dRTs = 23.71 ms) was significantly higher than in the other two experiments (all p <. 05; dRTs Exp.1 = 4.26 ms, dRTs Exp.2 = 14 ms). No significant difference was found between Experiments 1 and 2 (p = .09). As regards with Number-to-Space congruency, the ANOVA highlighted a significant main effect of Experiment [F (2, 81)

= 6.22, p

<

.01, η_p_^2^ = .14]. Post-hoc tests showed that in Exp. 3 congruency (dRTs = 24.35 ms) was higher than in the other two experiments (dRTs Exp.1 = .96 ms, dRTs Exp.2 = 12.66 ms; all p <. 05). No significant difference was found between Experiments 1 and 2 (p = .09).

*1.5.2 Reliability and variability testing.*

The reliability of Space-to-Number congruency effect observed in Exp. 3 (.85) was higher than those of Exp.1 (.39) and Exp.2 (.41) [F (2, 81)

= 1020.39, p

<

.001, η_p_^2^ = .96; all post-hoc comparisons p < .001]. In a similar way, the reliability of the Number-to-Space congruency effect found in Exp. 3 (.78) was higher than that of the effects found in Exp.1 (.56) and Exp.2 (.43) [F (2, 81)

= 478.76, p

<

.001, η_p_^2^ = .92; all post hoc comparisons p < .001].
Both in the case of Space-to-Number [F (2, 81)

= .40, p

=

.67, η_p_^2^ = .01] and Number-to-Space [F (2, 81)

= .71, p

=

.49, η_p_^2^ = .02] congruency effects, no difference in the variability of reliability measurements was found among the three experiments.
